# Preliminary Development of a Robotic Hip-Knee Exoskeleton with 3D-Printed Backdrivable Actuators

**DOI:** 10.1101/2023.05.18.541377

**Authors:** Alyssia Sanchez, Trent Rossos, Alex Mihailidis, Brokoslaw Laschowski

## Abstract

Robotic exoskeletons can provide powered locomotor assistance and rehabilitation to persons with mobility impairments due to aging and/or physical disabilities. Here we present the preliminary development and systems integration of T-BLUE - a modular, bilateral robotic hip-knee exoskeleton with 3D-printed backdriveable actuators. We retrofitted commercially available passive postoperative orthoses with open-source 3D-printed actuators to minimize cost and improve accessibility. The actuators are of quasi-direct drive design with high-torque density brushless DC motors and low gearing (15:1 transmission ratio) for low output impedance and high backdrivability, therein allowing for energy-efficient and dynamic human-robot physical interaction and legged locomotion. The modular design allows the exoskeleton to be customized and adapted to different users (e.g., persons with lateral vs. bilateral mobility impairments) and different hip-knee joint configurations. The goals of this preliminary study were to describe our experience with regards to the repeatability of the open-source 3D-printed actuators in engineering practice and the feasibility of integrating the actuators into wearable robotics hardware. This qualitative research serves as a first step towards using the robotic exoskeleton to support the development and testing of novel controller designs and rehabilitation protocols for different locomotor activities of daily living. We are especially interested in populations that could benefit from partial locomotor assistance such as older adults and/or persons with osteoarthritis. Future research will involve benchtop testing to quantitatively evaluate the actuator performance in terms of dynamics and energy-efficiency.

## 1. INTRODUCTION

Robotic exoskeletons are wearable assistive and rehabilitative devices that can provide physical training and enhance human mobility [1]. However, most commercial exoskeletons have been designed to provide high amounts of assistance to persons with complete paralysis [2],[3]. For example, the ReWalk device is the first powered exoskeleton to attain FDA clearance for personal and rehabilitation use and provides hip and knee motion to individuals with spinal cord injury. The system is an external orthotic design that actively controls the gait pattern of the legs and enables walking, sit-to-stand, and stand-to-sit movements [4]. Ekso Bionics developed the Ekso exoskeleton that bilaterally actuates the knee and hip joints and provides assistive walking. Another notable development is the Indego exoskeleton, which is marketed largely to persons with spinal cord injuries [4].

These commercial exoskeletons are often designed without leveraging the passive dynamics of human locomotion [5]. Due to the high torque requirements of the legs during walking and other daily locomotor activities, past exoskeleton designs have increased torque density by using high-ratio transmissions with low-torque high-speed motors, causing high output impedance [6]-[8]. These designs also lack backdrivability, which inhibit compliant and dynamic human-robot physical interactions. This can result in heavy, inefficient, and rigid actuators that require significant energy consumption and have limited battery-powered operation, which reduces their untethered use in real-world environments [9]. There is a large and growing population of persons who could benefit from partial locomotor assistance, such as older adults and/or persons with physical disabilities due to musculoskeletal conditions (e.g., osteoarthritis) and neurological disorders such as stroke.

Series elastic actuators have been used in legged and wearable robotic systems to help address some of these design challenges and improve torque fidelity, store energy, and reduce impact forces [10]. Another emerging design is quasi-direct drive actuators, which can mimic the elasticity of series elastic actuators by impedance control and reduce impact forces via lowering the output impedance and allowing for backdriving the system [10],[11]. Interestingly, the efficiency of quasi-direct drives can be bidirectionally asymmetric [12]. This use of high-torque motors with low transmission ratios are common in autonomous drones [13]. However, exterior rotor brushless DC motors used in drones often implement open-loop communication systems or electronic speed controllers, which may struggle to operate at the lower speeds characteristic of wearable robotic devices [14].

The MIT Cheetah robot used quasi-direct drives and low inertia legs for energy-efficient and dynamic locomotion [15]-[18]. The robot has a cost of transport of 0.5, which is significantly lower than other legged robots and can control contact forces during dynamic bounding, with contact times of 85 ms and peak forces greater than 450 N [18]. The MIT Cheetah demonstrated the benefits of quasi-direct drive actuators in legged robotics with promising future applications to wearable robotic systems for human walking. Because these actuators can provide low output impedance and high backdrivability, they can have many benefits over existing robotic prosthetic legs and exoskeletons, including freely swinging dynamic joint motion, compliance with the ground, and power regeneration [19]-[21]. For example, the robotic leg prosthesis by Elery et al. [22],[23] achieved high torque output (>180 Nm) despite using low-impedance actuators with minimal gearing.

Here we present the preliminary development and systems integration of T-BLUE - a modular, bilateral robotic hip-knee exoskeleton that can be used for locomotor assistance and rehabilitation. The design builds on open-source technologies, including retrofitted passive postoperative orthoses and 3D-printed quasi-direct drive backdriveable actuators from the University of Michigan, in order to create an accessible solution for energy-efficient and dynamic legged locomotion and human-robot interaction. The goals of this preliminary study were to describe our practical experience with regards to the repeatability of the open-source 3D-printed actuators and the feasibility of integrating the actuators into wearable robotics hardware. This qualitative research provides as a first step towards using the T-BLUE robotic exoskeleton to support the development and testing of novel controller designs and rehabilitation protocols.

## 2. ROBOTIC EXOSKELETON

### 2.1 Materials

The materials for the exoskeleton design relating to the mechanical brace included the bilateral Breg T-Scope Premier postoperative hip and knee orthoses and aluminum support struts to replace the existing struts in the orthoses (see Figure 1). We used waterjet machining to cut 7075 aluminum to create the support struts. 7075 aluminum was selected due to its high strength and resistance to stress and strain, which can be useful for repeated mechanical loading such as during daily lower-limb mobility and walking. To help minimize cost, the actuator design and mechanical brace support struts were obtained from open-source libraries [2]. The mechanical brace was modified using SolidWorks to interface with the 3D-printed actuators such that the circular attachment point of the support strut aligns with the actuators and the screw holes on the actuators while remaining suitable to fit the knee and hip orthoses.

**FIGURE 1.**
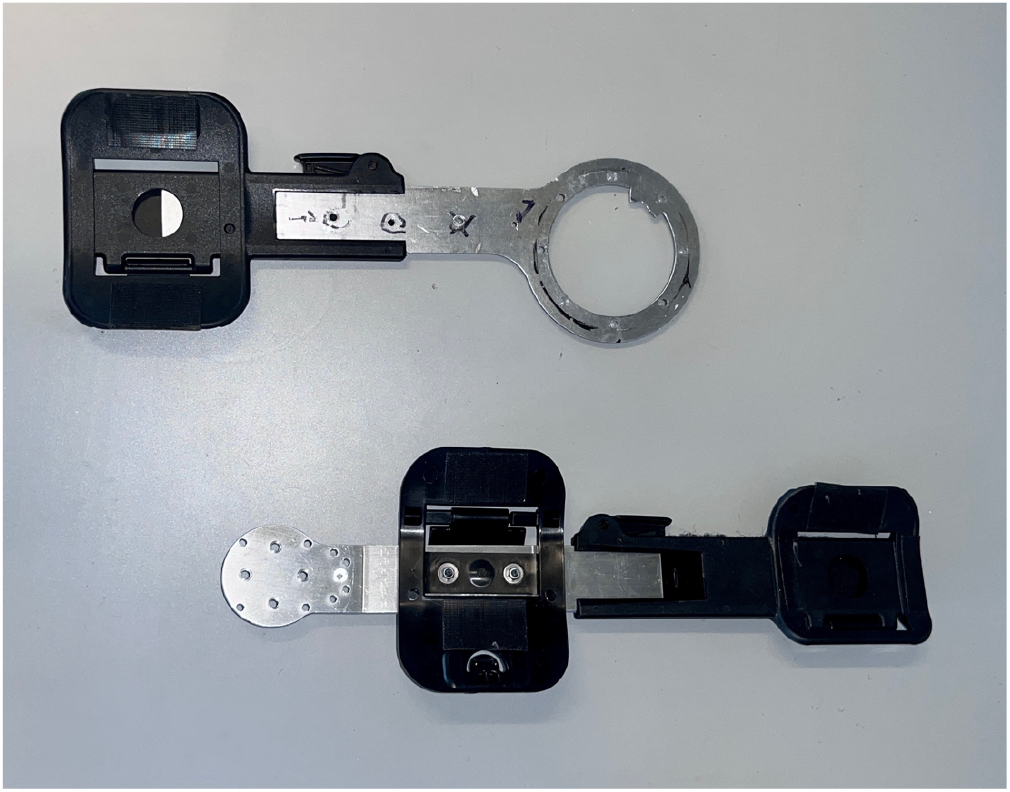
THE UPPER AND LOWER STRUTS OF THE EXOSKELETON MECHANICAL DESIGN, INCLUDING THE MACHINED METAL PARTS AND THE PASSIVE ORTHOSIS PADDING SUPPORTS FOR THE HUMAN-EXOSKELETON INTERFACE. PHOTOGRAPH OF KNEE JOINT.

The external components of the actuators were printed with ASA, a cost-effective and customary material used in 3D-printing. The components placed near the motor (i.e., the sun gear and motor housing) were printed with ULTEM 1010 resin, which has a higher melting temperature and is more robust than ASA, in order to ensure that heat emitted from the motor would not interfere with the performance of the actuator and to mitigate degradation risks. These 3D-printed materials differ from previous 3D-printed actuators for legged robotics [10], which used FDM-PLA and SLA-HT. The actuator components were printed using a Fortus 450MC 3D-printer with an accuracy of ± .127 mm (± .005 in.).

### 2.2 Actuators

We developed the T-BLUE exoskeleton using 3D-printed quasi-direct drive actuators originally designed and made open-source by the University of Michigan [10],[11] for reproducibility and design flexibility. The quasi-direct drive actuators cost less than $200 each and can allow for high-speed, responsive locomotion and achieve backdrivability and low inertia [10]. The bilateral drive is a compound-planetary transmission known as Wolfrom. To improve efficiency, the gear geometry had a 15:1 transmission ratio and to increase strength, all gears used a 30° pressure angle profile [10].

Figure 2 shows the inner planetary gear assembly. We press-fitted the dowel pins with the bearings to allow rotation of the three planetary gears within the assembly. This was one of the most complicated tasks as it required additional machine shop tools and careful assembly to ensure that the bearings were not damaged, and that the gears rotated smoothly. It was difficult to push the dowel pins into the bearings and gear carrier holes since the dowel pins had slightly greater tolerance (0.004 to 0.012 mm) than the bearings and holes in the 3D-printed parts. We eventually outsourced this process to a machine shop that used a hydraulic press. The dowel pins were sanded to fit into the upper and lower halves of the planetary gear assembly carrier. Prior to outsourcing this process to a machine shop, we tried manually pressing the pins into the bearings, and the bearings into the gears, which damaged the bearings and impacted the smooth actuation. Once the parts were pressed, bolts and nuts were used to secure the upper and lower carrier halves to secure the planetary gear assembly.

**FIGURE 2.**
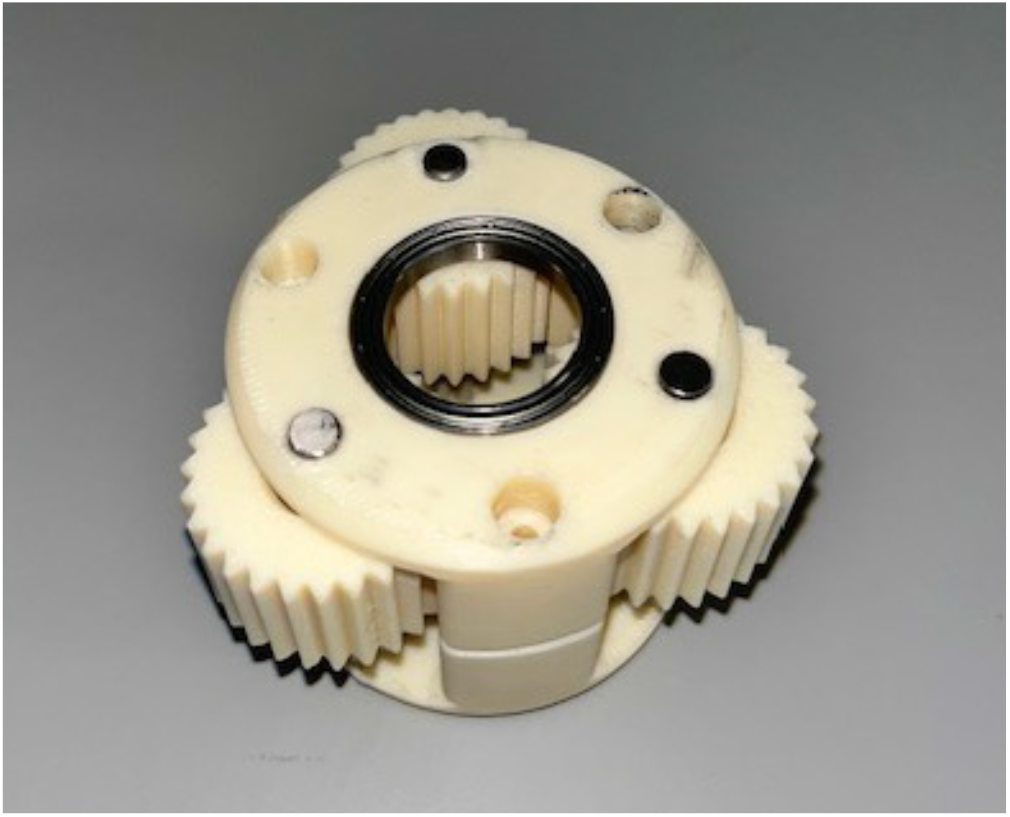
THE INNER PLANETARY GEAR ASSEMBLY OF THE 3D-PRINTED ACTUATOR WITH 3 PLANETARY GEARS, DOWEL PINS, AND BEARINGS, HELD TOGETHER BY THE UPPER AND LOWER CARRIER HALVES.

After building the inner planetary gear assembly, the outer encasing of the actuator was prepared. Figure 3 shows the individual grouped assemblies within the actuator, including (1) the inner planetary gear assembly, (2) the upper inner encasing, (4) the upper outer encasing that holds (3) the ring gear, and (5) the motor assembly. The ring gear is placed within the upper outer encasing, which meshes with the larger gear ends of the three planetary gears. We sanded the exterior circumference of the sun gear and press-fitted a R150 rotor magnet into the sun gear (Figure 4) and bearings were attached.

**FIGURE 3.**
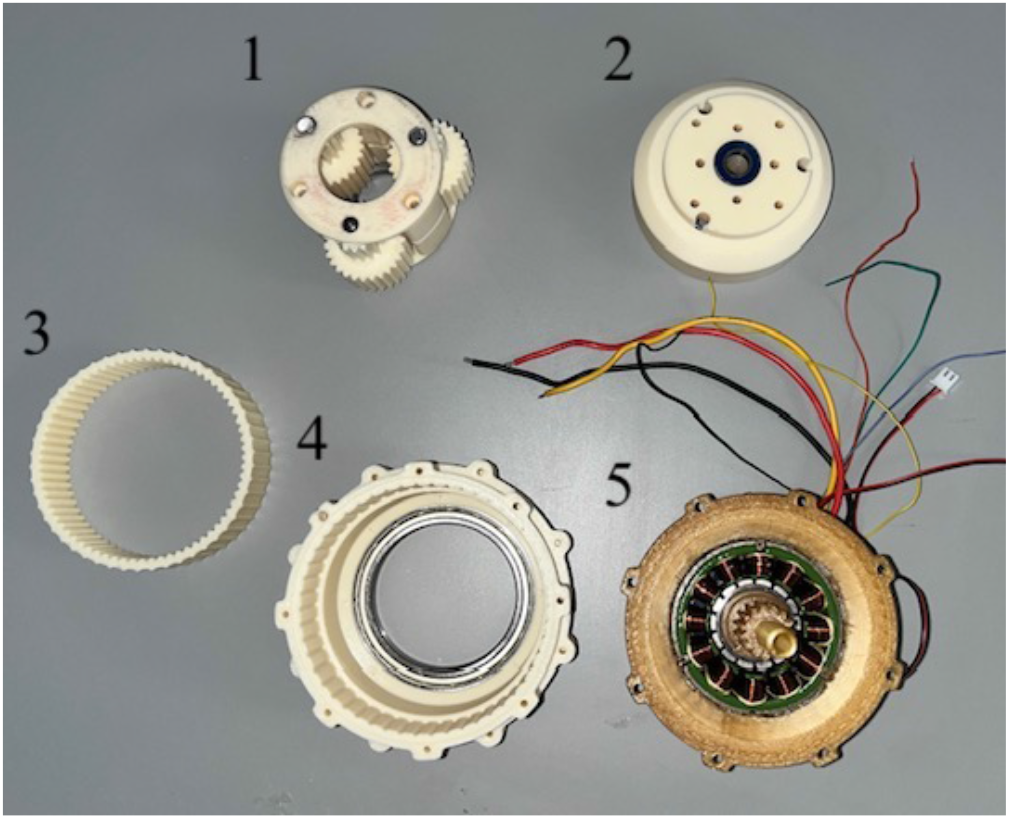
THE INDIVIDUAL GROUPED ASSEMBLIES WITHIN THE 3D-PRINTED ACTUATOR, INCLUDING (1) THE INNER PLANETARY GEAR ASSEMBLY, (2) THE UPPER INNER ENCASING THAT HOLDS THE INNER PLANETARY GEAR ASSEMBLY, (4) THE UPPER OUTER ENCASING THAT HOLDS (3) THE RING GEAR, AND (5) MOTOR ASSEMBLY.

**FIGURE 4.**
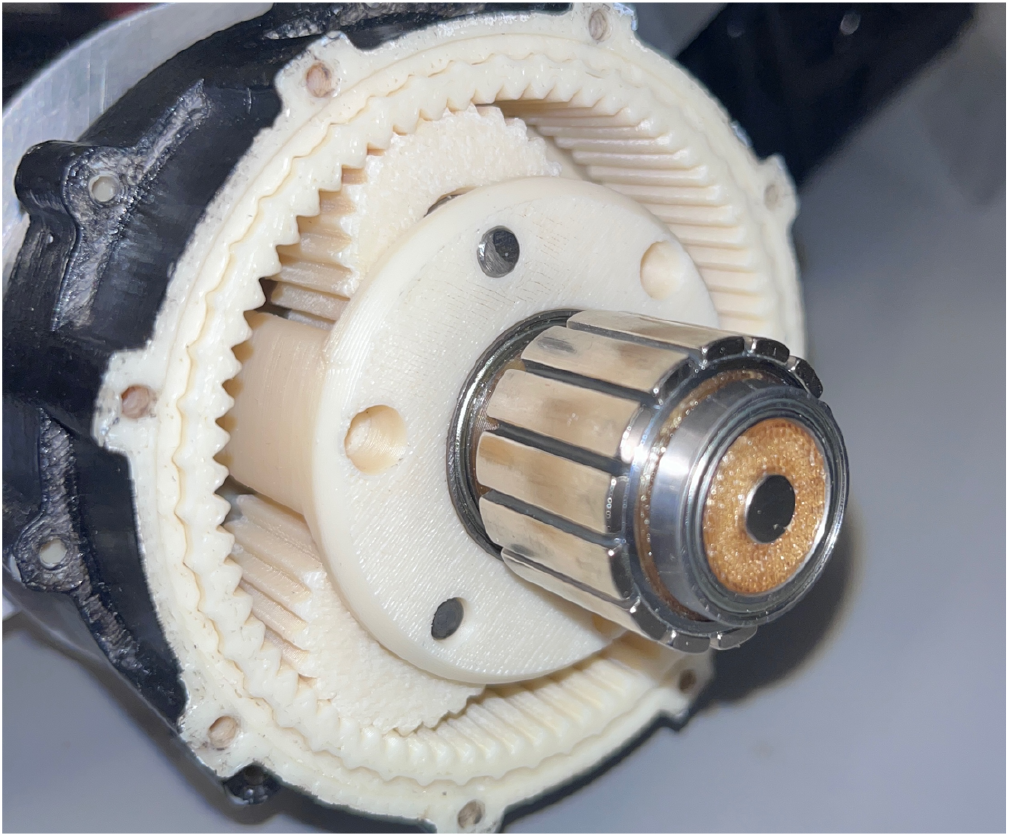
THE 3D-PRINTED ACTUATOR SHOWING THE ROTOR MAGNET ATTACHED TO THE SUN GEAR AND THE BEARING AT THE DISTAL END. THE SUN GEAR IS MESHED WITH THE INNER PLANETARY GEAR ASSEMBLY.

The motor encasing printed with ULTEM 1010 resin initially had leftover 3D-print support material that obstructed the openings and prevented attachment of the motor stator. Unlike the ASA 3D-printed parts that used soluble support material, the ULTEM 1010 resin used breakaway support, which was difficult to remove. We used box cutters to scrape away the excess 3D-print support. The sun gear with the R150 rotor magnet was pressed into the stator. This sun gear assembly was then pressed flush against the top of the upper outer encasing while ensuring no damage to the motor encoder. A Moteus board was used to control the motor (see Figure 5) and fans were attached to the motor encasing assembly to prevent overheating. We then integrated the different grouped actuator assemblies. This required pressing the assemblies together and using screws along the outer circumference of the actuator to secure the upper outer encasing to the motor assembly. The exterior of the actuators was painted with acrylic paint.

**FIGURE 5.**
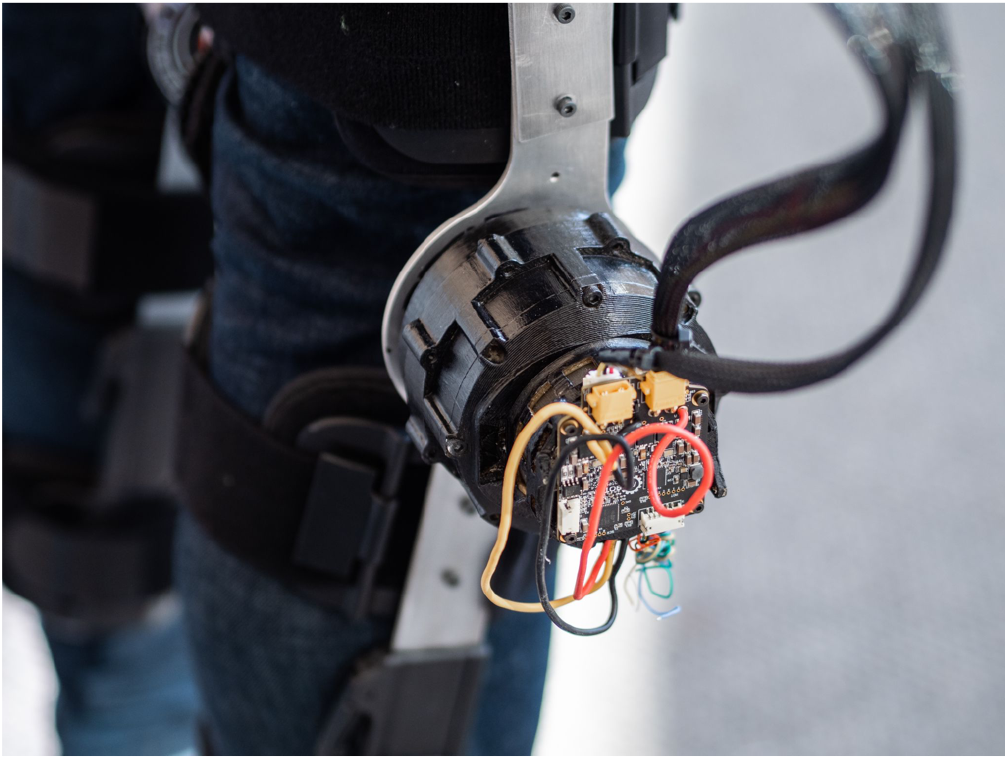
EXOSKELETON KNEE ACTUATOR WITH THE BRUSHLESS DC MOTOR CONTROLLER AND ATTACHED TO THE MACHINED METAL STRUTS.

While benchtop testing of our actuators is still ongoing, similar 3D-printed actuators [10] have shown thermal performance limits similar to metallic designs and high durability from testing over 420,000 strides of simulated walking. However, these 3D-printed actuators used FDM-PLA and SLA-HT materials, while we used ASA and ULTEM 1010 resin, respectively [10]. We selected the T-Motor R150 brushless DC motors based on a balance of cost, torque constant, motor constant, and inertia. Previous work [10] determined that, when the T-Motor R150 was paired with the bilateral drive 15:1, the ratios were appropriate for 8-15 kg legged robots, which we translated to wearable robotic systems. The open-source 3D-printed actuators can generate up to 38.2 Nm of peak torque and 8.8 Nm of continuous torque [10], which is comparable to lower-limb exoskeletons.

### 2.3 Electrical System

We used Mjbots Moteus r4.8 brushless motor controllers with magnetic absolute encoders, and Mjbots pi3hat, to control the motors via CAN technology. The pi3hat allows a Raspberry Pi 3b+ to communicate with the motor controllers over a CAN interface. The CAN bus simplifies the wiring path since each motor controller can be daisy chained. Two 5-cell 3300 mAh lithium polymer batteries were connected in parallel to power the exoskeleton. We used the Mjbots distribution board for safe and easy power distribution to each motor controller, which also provides real-time telemetry data to facilitate debugging. Although we currently have only one inertial measurement unit (IMU) integrated into the pi3hat, future work will include the integration of two IMUs into the design controlled by the Raspberry Pi via an inter-integrated circuit (I2C) communication protocol. The python package *moteus_gui* helped with configuring and debugging the actuator using real-time telemetry and provided a graphical user interface to edit motor configurations. To program the controller, we used python packages *moteus* and *moteus-pi3hat*. See [24] for full documentation of these libraries.

### 2.4 Mechanical Design

The mechanical design of our exoskeleton builds on the M-BLUE design by the University of Michigan [2], which we customized for the larger diameter of the 3D-printed actuators. We retrofitted passive postoperative orthoses with our actuators by removing the outer support struts on the orthoses and replacing them with machined metal struts. The lateral sides of the knee orthoses were disassembled, and we separated the paddings and padding supports from the struts. The metal struts that we machined to interface with the 3D-printed actuators were then attached to the paddings and padding supports. This procedure was performed similarly for the hip. The actuators were then attached to the metal struts, lateral to the knee and hip joints. During the initial attachment, the external medial-facing metal bearings of the actuator were not flush with the 3D-printed portion, which caused the metal struts to not sit flush with the actuators due to the extra clearance. We created 3D-printed parts to fit between the actuator and metal struts, and the orthoses straps were attached. Photographs of the fully assembled robotic hip-knee exoskeleton are shown in Figure 6.

**FIGURE 6.**
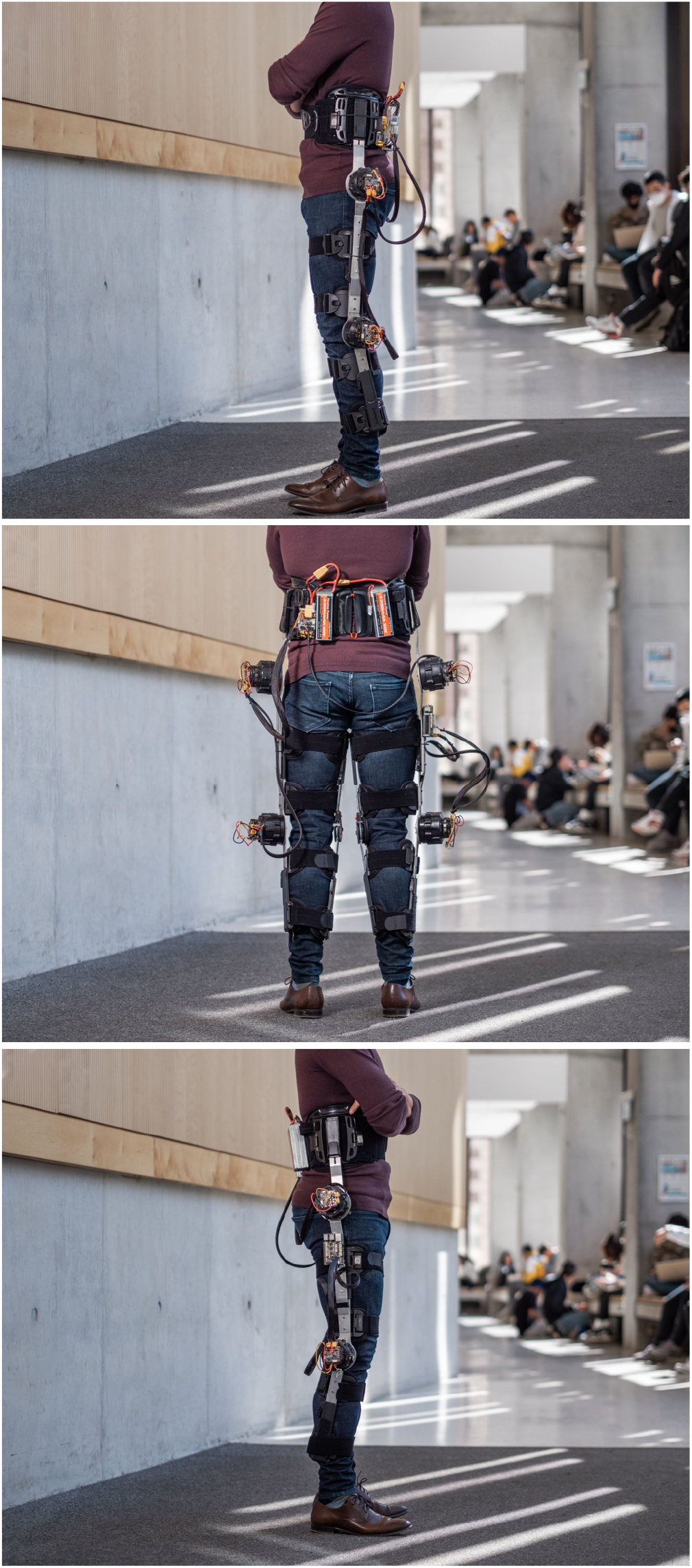
THE FULLY ASSEMBLED T-BLUE ROBOTIC HIP-KNEE EXOSKELETON, INCLUDING THE 3D-PRINTED QUASI-DIRECT DRIVE ACTUATORS, BRUSHLESS DC MOTOR CONTROLLER, MACHINED METAL STRUTS, AND RETROFITTED PASSIVE POSTOPERATIVE ORTHOSES.

## 3. DISCUSSION

In this paper, we present the preliminary development and systems integration of a modular, bilateral robotic hip-knee exoskeleton with 3D-printed backdriveable actuators, otherwise known as T-BLUE. Exoskeletons are wearable devices that can provide powered locomotor assistance and rehabilitation to persons with mobility impairments due to aging and/or physical disabilities. However, most exoskeletons use highly-geared rigid actuators, which have several limitations in terms of efficiency and control [4],[5]. In contrast, we used high-torque density brushless DC motors with low gearing (15:1 transmission ratio), also known as quasi-direct drives, to achieve low output impedance and high backdrivability, therein allowing for energy-efficient and compliant human-robot interaction and legged locomotion. To minimize cost and improve accessibility, we retrofitted commercially available passive postoperative orthoses with open-source 3D-printed actuators, originally designed by the University of Michigan [2],[10]. The goals of this study were to describe our practical experience with regards to the repeatability of the open-source 3D-printed actuators and the feasibility of integrating the actuators into wearable robotics hardware.

The robotic exoskeleton has several advantages compared to other exoskeletons. The modular design allows the robot to be customized and adapted to different users (e.g., individuals with lateral vs. bilateral impairments) and different hip-knee joint configurations. This adaptability can be particularly useful for persons who regain mobility in certain lower-limb joints at a faster rate than other joints [1]. Another advantage of the design is its simplicity. We leveraged commercial postoperative knee and hip orthoses, which allowed for easier material procurement, assembly, and donning and doffing. The exoskeleton support struts can be easily customized to users of different sizes since we designed them using computer aided design. Retrofitting commercial passive orthoses with the open-source 3D-printed actuators allowed us to develop the T-BLUE exoskeleton for roughly $5,100 CDN, including service charges for machining.

Unlike robotic exoskeletons that use highly-geared motor-transmission systems, which can have high energy consumption and peak power requirements [4],[5], T-BLUE is designed to exploit the passive dynamics of legged locomotion. The backdrivability of the actuators can allow for greater energy-efficiency and bidirectional power flow, including energy regeneration (i.e., the conversion of otherwise dissipated mechanical energy during periods of negative joint work into electrical energy) [19]-[21]. Simulations of human-exoskeleton walking have shown that actuators with energy regeneration can, in theory, extend the battery-powered operating times by up to 100% [21]. Moreover, the backdrivability and low impedance of the actuators can support volitional control (i.e., the person can actively move the joints), which can promote rehabilitation and user engagement. The low output impedance can also facilitate safe human-robot physical interactions.

When reproducing the open-source 3D-printed actuators, we experienced several unexpected challenges and limitations. For example, when building the inner planetary gear assembly, we initially damaged the metal balls inside the bearings by forcefully pushing the bearings into the gears, which caused the bearings to rotate with higher resistance as opposed to smoothly rotating. In some instances, this also resulted in damage to the gears. Fortunately, we had spare 3D-printed parts and extra bearings for replacement. We eventually outsourced the inner planetary gear assembly to a professional machine shop, which used a hydraulic press, to ensure a high-quality press-fit and to prevent further damage of materials.

Another challenge was ensuring that the actuators were assembled with a high degree of precision and consistency. We performed trial-and-error procedures as each actuator required careful assembly of the gears and precise placement of the bearings and the R150 rotor magnet. If the bearings or rotor magnet were not properly pressed, the upper and lower outer encasings of the actuator would not press tightly together, and thus prevent assembly of the actuator. Difficulties in assembling the actuators were largely due to the small, geometric tolerances in the hardware and 3D-printed actuator parts. As mentioned in Section 2.1, the accuracy of our 3D-printer is ± .127 mm (± .005 in.), while the dowel pins in the inner planetary gear assembly have a geometric tolerance of 0.004 to 0.012 mm. We were able to physically observe these seemingly small tolerances in the slight differences in the actuation smoothness between the four actuators that we built.

Another challenge was ensuring that the threading of the screw holes in the 3D-printed actuators remained intact during assembly. Each screw hole was manually threaded using steel tap to properly secure the screws. However, the ASA and ULTEM 1010 resin material is not ideal for repetitively inserting and removing screws, as the fibers within the screw holes eventually deteriorate and become stripped. To prevent mechanical wear of the 3D-printed material, metal inserts could be used in future design iterations. There can also be issues with mechanical slippage during actuation. If threading wear occurs, the actuator and housing no longer move as one rigid body, which can lead to further mechanical inefficiencies during power transmission.

Future research with the 3D-printed actuators will include benchtop experiments for lifecycle testing to evaluate durability and system identification to design physics-based computer models. As discussed in Section 2.2, [10] showed minimal performance degradation of the actuators over 57 hours of testing, or approximately 420,000 strides of simulated gait. However, given that different 3D-printed materials will have different strengths, the ASA and ULTEM 1010 resin material that we used in our actuators will have, to some extent, different durability results than the FDM-PLA and SLA-HT materials used by [10]. The benchtop testing will also serve to characterize the active and passive dynamic behaviors of the actuators in order to design computer models for simulation and optimization. This builds on some of our previous dynamometer testing with robotic exoskeleton actuators [21].

The long-term purpose of the T-BLUE exoskeleton is to support the development and testing of new controller designs and rehabilitation protocols for different activities of daily living (e.g., walking, stand-to-sit, sit-to-stand, and ramp and stair ascent and descent). We are especially interested in populations who could benefit from partial locomotor assistance such as older adults and/or persons with osteoarthritis or other musculoskeletal injuries. This differs from most robotic exoskeletons, which have been designed to provide complete assistance to persons with spinal cord injuries [4]. In addition to movement assistance, exoskeletons can promote rehabilitation while simultaneously performing activities of daily living, as well as human performance augmentation (e.g., providing caregivers additional strength when tending to patients). These relatively untapped applications using wearable robotic systems like the T-BLUE exoskeleton motivate our future work.

## ACKNOWLEDGEMENTS

We acknowledge and thank the University of Michigan for the original exoskeleton design and open-source 3D-printed actuators. We also thank Deniz Jafari, Hannah Smegal, and the University of Toronto Department of Mechanical and Industrial Engineering machine shop for helping the exoskeleton assembly. We dedicate this research to the people of Ukraine in response to the 2022 Russian invasion. This work was supported by The Schroeder Foundation.

